# Antigen density and applied force control enrichment of nanobody-expressing yeast cells in microfluidics

**DOI:** 10.1101/2024.01.03.574015

**Authors:** Merlin Sanicas, Rémy Torro, Laurent Limozin, Patrick Chames

**Affiliations:** Aix-Marseille Université, CNRS, INSERM, Institute Paoli-Calmettes, CRCM, Marseille, France; Aix-Marseille Université, CNRS, INSERM, LAI, Marseille, France

**Keywords:** Yeast Display, Laminar Flow Chamber, nanobodies, Cell Enrichment

## Abstract

In vitro display technologies such as yeast display have been instrumental in developing the selection of new antibodies, antibody fragments or nanobodies that bind to a specific target, with affinity towards the target being the main factor that influences selection outcome. However, the roles of mechanical forces are being increasingly recognized as a crucial factor in the regulation and activation of effector cell function. It would thus be of interest to isolate binders behaving optimally under the influence of mechanical forces. We developed a microfluidic assay allowing the selection of yeast displaying nanobodies through antigen-specific immobilization on a surface under controlled hydrodynamic flow. This approach enabled enrichment of model yeast mixtures using tunable antigen density and applied force. This new force-based selection method opens the possibility of selecting binders by relying on both their affinity and force resistance, with implications for the design of more efficient immunotherapeutics.

## Introduction

Immune cells apply and sense for mechanical forces that aid in cellular motility and in probing their proximal environment. Lymphocytes in particular have several modes of motility that makes use of mechanical forces depending on the environment that they are traversing: an integrin-dependent motility, an amoeboid-like adhesion-independent motility and the rolling, adhesion and transmigration used in long distance travel through blood vessels (1–3). Activating receptors of lymphocytes have also been shown to both apply and sense mechanical forces. T cell receptors (TCR) and their interaction with the peptide major histocompatibility complexes (pMHC) have been studied intensively for the past decades. In order to probe their environment, T cells generate piconewton (pN) forces to the pMHC (4,5) which may be involved in peptide discrimination (1). Indeed, the discrimination capabilities of the TCR is now admitted to be encoded in the life-time distribution of TCR-pMHC bonds, while equilibrium affinity is not sufficient to explain the exquisite capacity of the T cell to find rare agonists in a sea of non-agonists (6). The bond lifetime is modulated by the force applied to the bond, leading to catch bonds exhibiting longer lifetime under force or slip bonds exhibiting the opposite behaviour. Those behaviour have been shown to play a central role in TCR-pMHC recognition (7–10). For Natural killer (NK) cells, the interaction of activating receptor NKG2D with one of its ligands, MICA, has been shown to be mechanosensitive (11) and is hypothesized to have catch-bond properties (12). Likewise, B cells physically pull on their target antigens to differentiate between a high affinity and a low affinity antigen (1,13,14), though the existence of a catch-bond BCR or antibody has yet to be clearly established (15). To our knowledge, only nanobodies (Nbs), corresponding to the variable fragment of the Heavy-chain only antibodies (VHH) from the Camelidae serum (16) have been shown to have catch bond properties, as shown for a Nb binding to FcγRIII (CD16) (17). This result suggests that Nbs could be selected to deliver biophysical cues leading to optimal immune cell activation and function.

Biophysical methodologies can be used to measure the force dependence of these ligand-receptor interactions at the cellular or molecular level. Single interaction techniques include Biomembrane Force Probe (BFP) (1), Optical Tweezers (1) and Atomic Force Microscopy (AFM) (18). Options with higher throughput include magnetic tweezers (19), acoustic force spectroscopy (AFS) (20,21) or the laminar flow chamber (LFC) (28). LFC uses microbeads coated with a specific receptor driven along the surface of a small channel derivatized with a very diluted cognate ligand. The interaction is viewed through a microscope focusing on the bond formation and rupture resulting in transient bead arrests under flow (17,22–24). In conditions of single bond observation, a direct measure of bond lifetime under force can be obtained. Similar microfluidic devices were used to immobilize target cells by coating the microfluidic surface with a capture antibody and flowing cells that present the cognate antigen on their surface (25–27).

In vitro display technologies have been versatile and powerful tools for the discovery of proteins that bind specifically to a target. This started with phage display (28) which paved the way to other display methods such as yeast display (29). In this case, the protein of interest is expressed on the yeast surface fused genetically to an anchor protein. Whole antibodies or antibody fragments such as single chain variable fragments (scFv) or Nbs can be expressed on the yeast surface. For instance, yeast display has been used in the discovery of scFvs that bind to West Nile virus envelope protein (30), antibodies against Botulinum neurotoxins (31), and Nbs that target human GPCRs (32) or SARS-CoV-2 receptor binding domain (RBD) (33). Advantages of yeast display include compatibility with flow cytometry, ease of manipulation and handling of yeast cells, eukaryotic post translational modifications and proper folding of the expressed proteins (34). A direct comparison between yeast and phage display using the same cDNA library of scFv showed that yeast display isolated more unique binders compared to its phage display (35). However, when used for antibody or antibody fragment selection, yeast display, similar to other in vitro display technologies, relies on antigen-antibody interaction in solution and is dictated by affinity alone, with no consideration to force sensitivity of the interaction.

In the past few years, several studies combined microfluidics and mycology, such as the so called ‘Fungi-on-a-chip’ platforms (36), one of which was used for adhesion-based cell separation (37). Here, we present a novel assay that combines Nb yeast display and LFC to capture yeast cells under flow in an antigen-specific manner. Two microfluidic devices were used, one to measure the antigen specific and non-specific adhesion of Nb-expressing yeasts, and another for enrichment of model mixtures of Nb-expressing yeasts analyzed by imaging and cytometry. The assay directly quantifies the adhesive properties of two different Nb-expressing yeast strains by monitoring the number of cells captured before and after flow. Force applied on cells was controlled through the shear rate to induce detachment of non-specifically adhering cells from the surface while maintaining antigen-specific adhesion. Furthermore, we demonstrate how this device can be used for the enrichment of yeast displaying a antigen specific Nb under controlled antigen density and applied force, which has implications for the selection of Nbs with high affinity and resistance to force.

## Methodology

### Design of nanobody-expressing Yeast

The plasmid pYDS containing the pGAL1 for Nb expression, α-mating factor leader sequence, HA tag and 649 stalk sequence (32) was modified to contain either the Nef19 Nb (38) or CD16.21 Nb (17,39) gene using HiFi DNA Assembly Cloning kit (E5520S, New England BioLabs Inc.). The plasmids were transformed via LiAc/SS-carrier DNA/PEG method (40) into *Saccharomyces cerevisiae* (BJ5465).

### Cytometry of yeast

For cytometry, 2 x 10^6^ induced yeast cells were pipetted into wells of a V-bottom 96 well cytometry plate. The plate was centrifuged at 3500 x g, 4 °C for 1 min and the pellets resuspend with 100 µl PBS 1x with 0.2 % BSA; this was repeated 3 times. The pellets were resuspended in 100 µl mixture of Nef-ATTO 647N or CD16a-ATTO 647N (10 nM) with anti-Hemagglutinin-PE (aHA-PE, 0.375 µg/mL, Clone GG8-1F3.3.1, 130-120-717, Miltenyi Biotec) in PBS 1x with 0.2 % BSA and incubated at 4 °C on a platform shaker for 1 h. After, the plate was washed 3x and fixed using PBS 1x containing 0.2 % BSA and 1 % Paraformaldehyde (PFA) diluted from 16 % PFA (043368.9M, Thermo Scientific). This was incubated at 4°C for 15 min on a platform shaker. After 3 washes using PBS 1x. MACS Quant (Miltenyi Biotec) cytometer was used to perform flow cytometry experiments. Cytometry channel settings used were as follows: Forward Scatter (FSC): 300 V, Side Scatter (SSC): 420 V, B1 (CFSE): 260 V, B2 (PE): 290V, V1 (Alexa Fluor 405): 240 V all on hlog. R1 (ATTO 647N) settings were measured at two different values, 440 and 580 V, to adjust to signal differences on the monoclonal yeasts. Compensation settings were as follows: V1 at VioBlue 1, B1 at FITC 1, B2 at VioBlue 0.01, B2 at PE 1, R1 at APC 1. Trigger setting was at FSC: 1 and Events: 30,000. Analysis of flow cytometry data was done using FlowLogic 8.6 (Inivai Technologies Pty Ltd) and apparent affinity was estimated using Prism v5.03 using the non-linear regression function log(agonist) vs response – variable slope (four parameters).

### Growth and preparation of yeast

The transformed yeast cells were grown as described in previous literature (32). Tryptophan drop-out media (-Trp) was used for cell culturing. Solid -Trp plates (3.8g Tryptophan drop-out media supplement, 6.7 g Yeast Nitrogen Base, 20 mL Penicillin-Streptomycin, 2 % v/v glucose, pH 6) were prepared with 1 L to 12 grams agar ratio. Liquid -Trp with glucose medium had the same compositions as solid -Trp medium except agar. Liquid -Trp with galactose medium also had the same composition as the previous liquid medium except for switching 2 % v/v glucose to 2 % v/v galactose.

For cell culturing, yeast cells were first grown on solid -Trp plates. To prepare cell suspension, a single colony of yeast growing on a designated -Trp plate was scraped up with a sterile inoculating loop and suspended in 10 mL liquid -Trp with glucose medium. Suspended yeast cells were incubated in an Erlenmeyer flask at 30 ℃ with shaking at 220 rpm for 24 h. The suspension was centrifuged at 3500 x g for 1 min at room temperature. The pellet was resuspended in 10 mL -Trp with galactose medium and incubated in a new Erlenmeyer flask at 25 °C with shaking at 220 rpm for another 24 h to induce Nb expression on yeast surface. Cells were then prepared at an OD_600nm_ = 1 (1.5 – 3.0 x 10^7^ cells/mL) in -Trp with glucose medium.

### Fabrication of Microfluidic Device

The 1 entry – 1 exit design was based on the design of the commercial µ-Slide VI 0.4 (80601, Ibidi) while the 2 entries – 2 exits design was a modified version from another publication (37) both shown in Fig 2A & Fig 3A, respectively. Both devices were prepared as a three-layer sandwich. The thick top layer of Polydimethylsiloxane (PDMS) was prepared using SYLGARDTM 184 Silicone Elastomer kit at a 10:1 ratio (10 Liquid PDMS to 1 curing agent) and mixed thoroughly. Bubbles were removed by centrifugation at 1500 rpm for 2 min. After, the liquid was poured on a large 150 x 15 mm circular petri dish to reach a height of 6 mm and de-gassed during 30 min to remove bubbles. The petri dish was transferred to a 65°C oven to be cured for at least 3 h. The middle part was prepared by cutting a commercial 250 µm thin PDMS sheet (Sterne Silicone Performance) using a Graphtec Craft Robo Pro. The channel design and dimensions were transferred into the software of Graphtec, and the cutting was performed automatically after aligning the cutter. The lowermost portion, a standard 75 x 25 mm microscope glass slide (1.2 – 1.5 mm, Fisher 1239-3118), was washed with MilliQ water followed by 5 % Decon 90, MilliQ, 96 % Ethanol, MilliQ and Isopropanol and dried using nitrogen, followed by surface treatment using oxygen plasma (Harrick Plasma) at high setting for 10 min. The sides to be fused were placed in the chamber facing up. Simultaneous to the 10-min plasma treatment, the previously cut 250 µm thin PDMS was cleaned using MilliQ, Ethanol 96 %, MilliQ and 5 % Decon 90 and dried using nitrogen. Once dried and the microscope slide plasma treatment was finished, the clean 250 µm PDMS was also placed in the chamber alongside the microscope slide to be treated with oxygen plasma for only 2 min at low settings. Once finished, the thin PDMS layer and the glass slide were removed. The treated surfaces of each layer were apposed firmly afterwards and were placed on a hot plate at 95 °C for 10 min glass side bottom. The thick 6 mm PDMS layer was cut to a 75 x 25 mm dimension using a scalpel and the previously designed channel entry points were punched appropriately using a 4 mm diameter puncher. This thick PDMS was also cleaned and dried like that of the thin PDMS layer except for the portion of the 5 % Decon 90 where the punched PDMS was placed in a beaker and sonicated for 10 min. The thick PDMS and the glass-thin PDMS were once again treated with plasma oxygen at low settings for 2 min with the PDMS layers facing up. The treated surfaces were apposed firmly and again placed on the 95 °C hot plate for 10 min. The channels were placed in the 65°C oven for at least 3 h prior to use.

### Antigen Functionalization on Chamber Surface

Biotinylated BSA in PBS 1x (100 µg/mL, aliquoted from 10 mg stock, A8549-10MG, Sigma-Aldrich) was adsorbed directly on the channel surface and incubated for 1 h at room temperature on a tilting shaker. After 3 washes with PBS 1x, streptavidin in PBS 1x (10 µg/mL, 434302, Invitrogen) was incubated on the biotinylated BSA for 1 h. After 3 washes with PBS 1x, the chamber was incubated with Nef-biotin or CD16a-biotin in PBS 1x with 0.2 % BSA for 1 h. For the optimization experiment, a serial dilution was done with concentrations from 135 nM to 0.56 nM with 1/3 dilution factor per condition. For enrichment experiments, a constant concentration of 45 nM was used. The antigen incubation step was followed by 3 washes of PBS 1x followed by a passivation step with PBS 1x containing 2 % BSA to block the uncoated channel surface and incubated for 1 h. After 3 washes with PBS 1x, the channel was ready to be used in the LFC experiment.

### Microscope Settings

Microscopy was done using an inverted microscope (Axio Observer D1, Zeiss), controlled with Micro Manager 1.4.23 software and equipped with a 10x NA objective (Olympus A10PL 10x 0.25) with a 1.6x additional magnification. For fluorescence images, the light source used was PE-300 ultra (CoolLED) applying 100 % blue light (460 nm). For transmission images, halogen lamp at voltage 6 V was used. Fluorescence was recovered using Zeiss Filter set 16 (488016-9901-000, BP 485/20, FT 510, LP 515). Images were taken using Andor iXonEM + camera. Exposure times used were 10 or 500 ms for transmission microscopy images or fluorescence images, respectively). Electronic gain for fluorescence was set at 100. Pixel sizes of images corresponded to 0.787 µm per pixel. 8 images per condition (denoted after Preflow or Postflow) were taken at a 1000 µm distance lengthwise from the previous field of view. For experiments that included fluorescence images, the set of light microscopy images were taken first followed by going back to the initial field of view and manually switching to fluorescence imaging to take the same exact field of views.

### Assay for Capture Optimization

For capture optimization experiments, pure populations of Nef19^+^ or CD16.21^+^ yeast cells were used in the 1 entry-1 exit coated PDMS channels (Fig 2A). The device was connected using custom piping (Polytetrafluoroethylene (PTFE) Tube, 0.8 mm inner diameter x 1.2 outer diameter, PTFE Tube Shop), with a 3-way valve (Masterflex®, MFLX30600-25) to allow manual control between a 10 mL glass syringe (549-0539, VWR) mounted on a syringe pump (Pump 11 Pico Plus Elite, Harvard Apparatus) or an entry point for the yeast suspension. The exit pipe was directly placed over a beaker. Prior to beginning any experiments, the channels were purged with –Trp media with 2 % v/v glucose and Penicillin-Streptomycin to ensure that no bubbles were within the circuit. Induced yeast cells were prepared at a density of 7.5 x 10^6^ cells/mL and passed through a 27G 7/8-inch needle 10 times to dissociate yeast clumps (41). The yeasts sample was transferred to a sterile 1 mL plastic syringe and inserted on the appropriate 3-way valve entry port of the 1 entry-1 exit channel. The valves were adjusted to ensure that the direction of the yeast suspension was towards the channel. Once infused, a 5 min incubation period was given to allow majority of the yeasts to sediment to the surface and allow for Nb-antigen interaction. 1 min prior to the end of the incubation period, 8 bright field (BF) images were taken across the length of the channel (1 mm distance between each picture taken) and represents ‘PreFlow’ images. We calculated the shear rate (*G*) in 1/s applying the formula (42): *G = 6Q/ lh^2^*, using the channel width *l*, height *h* and flow *Q*.. Shear rates applied were varied between 168 and 926 1/s to apply a total volume of 6 mL per condition; the antigen incubation concentrations tested were 0.56, 1.67, 15, 45 and 135 nM. After the wash flow, another 8 pictures were taken again across the length of the channel and represents ‘PostFlow’ images.

### Assay for Enrichment

The 2 entries – 2 exits channels (Fig 3A) were used and a mixture containing 1:1, 1:10 and 1:100 binders to non-binders ratios of Nb-expressing yeasts were used. Each of the entry and exit was fitted with custom piping connected to a 3-way valve to allow the proper control of shear rate and flow direction during the entire enrichment process. This set-up required two different 10 mL glass syringe connected to the extreme ports (1st and 4th) and changed manually according to the needed direction of the flow. A schematic in Fig 3B illustrates the sequence of flowing and washing steps performed. First, both ports at the extreme ends (1st and 4th) were closed and the inner ports were opened (2nd and 3rd). The yeast cells were infused on the 2nd port, exiting to the 3rd port and allowed to sediment for 5 min. Pictures were taken as previously described in the optimization set-up, in bright field and in fluorescence (in the case of labelled negative yeasts). The 3-way valves were re-adjusted in such a way that the 1st port was closed, the 2nd port opened, the 3rd port closed and the 4th port opened. The 1st wash step was done with the direction of flow from the 4th port towards the 2nd port for 5 min at a shear rate of 337 1/s. The valves were again re-adjusted to have the 1st port opened, the 2nd port closed, the 3rd port opened and the 4th port closed. The 2nd wash step was done with the direction of flow from the 1st port towards the 3rd port, maintaining the shear rate and duration as wash 1. After the 2nd wash, pictures were taken representing ‘PostFlow’ images, in bright field and in fluorescence if relevant. The final adjustments of the ports were opening the extremes (1st and 4th) and closing the inner ports (2nd and 3rd). The elution phase, applying a significantly higher shear rate at 4800 1/s was used with the direction from 1st port to 4th port to detach the captured yeasts on the channel and recover them directly into a sterile 5 mL syringe attached to the 3-way valve in the 4th port and transferred into a 15 mL falcon tube. A 100 µl aliquot was recovered for cell counting. The recovered yeasts were concentrated into a 700 µl volume of –Trp with 2 % v/v glucose and Penicillin Streptomycin and incubated at 30 °C shaking at 220 rpm for at least 2 days in a 96-deep well plate. The media was changed into –Trp with 2 % v/v galactose and Penicillin Streptomycin and expanded to a volume of 5 mL in an Erlenmeyer flask and incubated for 1 day to induce expression. The induced yeast underwent cytometry to assess for enrichment.

### Yeast Cell Detection and Image Analysis

To process the captured images, FIJI (ImageJ 1.53t) was used with a specific script that employed the plugin MorphoLibJ to perform Gray Scale Attribute Filtering (Operation = Top Hat, Area minimum=100, connectivity=4), thresholding (1400), and particle detection using ‘Analyze Particles’ (size=4 – infinity pixels, circularity=0.1-1.00), an example of this is shown in Suppl. Mat. Fig 1. Applying the conversion factor of 0.787 µm per pixel, detection threshold was set at a minimum of 3.15 µm. The cell counts and other parameters were such as the centroid of every detected cell and the XY coordinates within the image were saved as csv files. The detected yeast cells were saved as regions of interest (ROIs). This macro was compared to a manual annotation as ground truth. We applied a machine learning program, TrackPy (43), for the Fluorescence imaging and matching using locate yeast function. Spot diameter was set to 11 pixels and adjusted at the minimum integrated brightness of the spot (minmass) to minimize detection of false positives as established on a sample of unstained yeast cells. For matching, we performed a linking of the cells detected on the bright field (MorphoLibJ generated csv files) and attributed them to the nearest fluorescent cell at a maximal distance of 50 pixels. The matched cells were temporarily removed and the process was reiterated to match some of the left-over cells that are still within the 50-pixel distance threshold. All the detections were reassembled and evaluated for the fraction of yeast cells that have found a fluorescent match.

## Results and Discussion

### Yeasts express functional nanobodies on their surface

Yeasts transformed with a vector bearing the GAL1-10 promoter and encoding surface expression of HA-tagged Nbs directed against HIV-1 Nef (Nb Nef19) or against human CD16 (Nb CD16.21) were expanded and induced using the presence of galactose in the –Trp medium. These yeasts were incubated with aHA-PE (0.375 µg/mL) to assess for the expression levels of the Nef19^+^ and CD16.21^+^ yeasts by flow cytometry. The average expression levels were 34 to 41 % (Fig 1A and 1B). For comparison, other publications that used the same plasmid reported expression levels of ∼25 % (32) and up to 70 %(44). We next studied the functionality of the expressed Nbs based on the binding schemes shown in Fig 1E.

**Figure 1.**
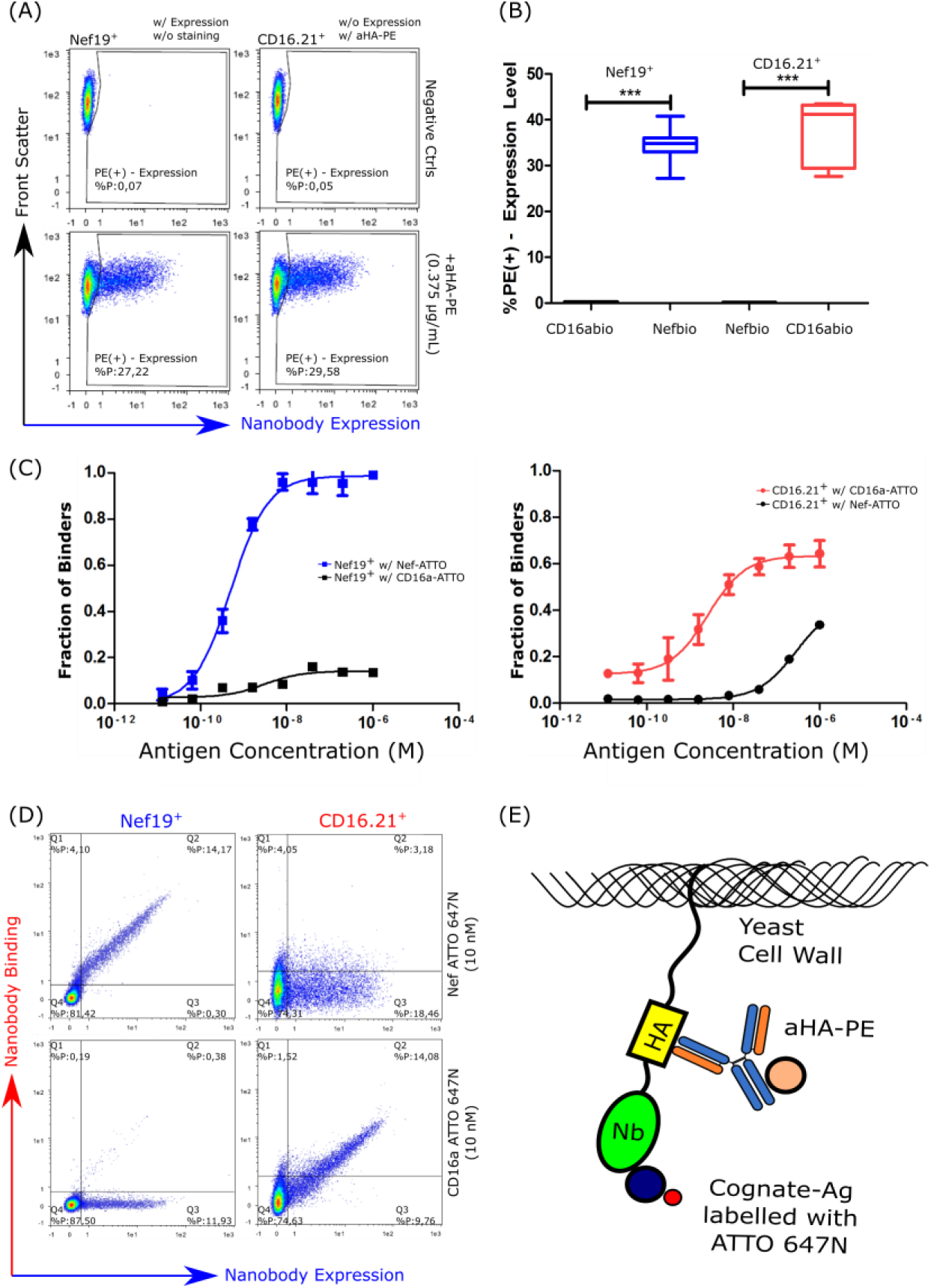
Nanobodies on the Yeast Surface. (A) Cytometry scatter plot of Nef19^+^ & CD16.21^+^ yeasts. (B) A vertical box & whisker plot corresponding to 3 independent cytometry measurements of the percentage of expressing yeasts using a-HA PE. Error bars are the standard error of means (SEM) (s = 3) (*** indicates a Student’s test with p ≤ 0.001). (C). An XY plot (fraction of binders as function of antigen concentration in M). The values on the x-axis were taken from a cytometry histogram elaborated in the Suppl. Mat Fig 2. This experiment was done 3 independent times. (D) 4 Quadrant gated scatter plots of the Nb-expressing yeast. The Nb-expressing yeasts were incubated with 10 nM of either antigen. (E) A schematic of the yeast cell wall when incubated with aHA-PE and their cognate antigen (Ag) with ATTO 647N.

To measure the apparent affinity of the Nef19^+^ and CD16.21^+^ on yeast surface, we performed a serial dilution experiment with their cognate or irrelevant antigen labelled with ATTO 647N, while fixing the concentration of a-HA PE. The measured apparent affinities were 5.4 x 10^-10^ M for Nef19^+^ and 2.6 x 10^-9^ M for CD16.21^+^ (Fig 1C). These findings were comparable to the previous publication result of 2 x10^-9^ M for Nef19 (38) and 1.0 x 10^-8^ to 1 x10^-9^ M for CD16.21 (17,39). An example of 4 quadrant gating of these yeasts in either their cognate or irrelevant antigen is shown in Fig 1D.

### Fluid-driven Yeasts specifically adhere to channel surface antigen

The Nb-expressing yeasts were driven along the surface of the 1 entry – 1 exit microfluidic channel derivatized with either their cognate or irrelevant antigen. The design of the microfluidic device and the application of flow and sequence of steps are shown in Fig 2A. The binding of the Nb-expressing yeast to their cognate antigens in flow is schematized in Suppl. Mat. Fig 3. The applied shear rates and antigen concentration on the channel surface were adjusted to maximize the ratio between specific and non-specific yeast capture using pure populations of Nef19^+^ or CD16.21^+^ yeasts separately. The surface of the channels were functionalized with various concentrations of target or irrelevant antigens, as performed for single bond measurements in LFC (17,22). We directly measured the effects of these changes by microscopy by counting the starting number of cells prior to flow and the remaining cells after the flow, as shown on bright field images of Fig 2B & 2C. The capture percentage (% Arrested Cells) was calculated by dividing the PostFlow count by PreFlow count and multiplied by a 100.

**Figure 2.**
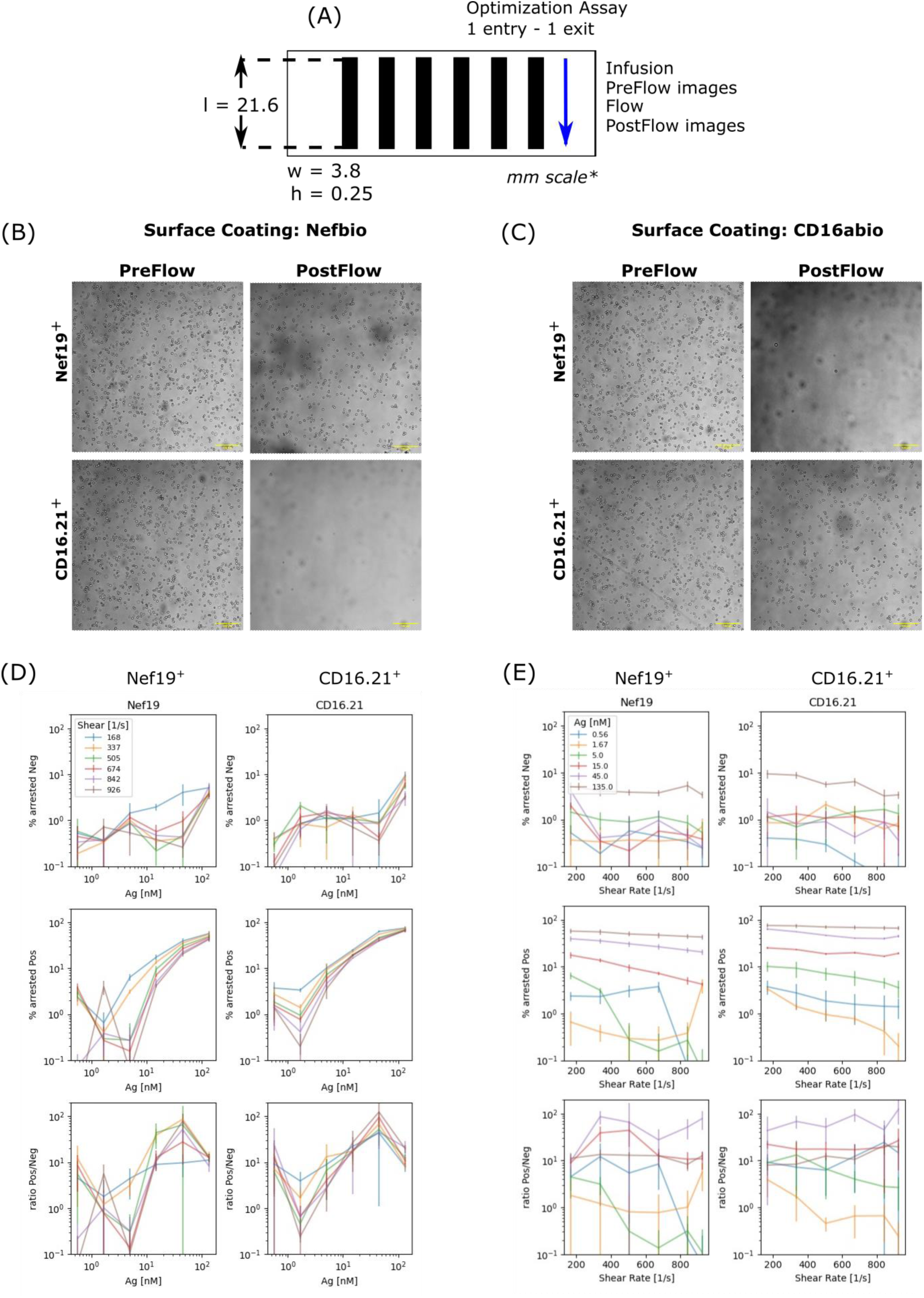
Optimization Assay microfluidic chamber, visualizing and optimizing specific adhesion by varying antigen concentration and Shear Rate. (A) 1 entry – 1 exit channel used for optimization based after the Ibidi µ-Slide VI 0.4. The direction of flow and indicated by the blue arrow and the sequence of steps are infusion of yeast cells, image acquisition prior to flow, application of flow and image acquisition after flow. The images of yeast cells driven along the surface of a channel incubated with either (B) Nefbio or (C) CD16abio. The top row corresponds to the Nef19^+^ yeast while the bottom row is for CD16.21^+^ yeast. The first and second column show a representative image PreFlow and PostFlow, respectively. (D) The columns correspond to the monoclonal yeast driven along the surface of a channel at varying concentrations of antigen. The 1^st^ row shows % Arrested on irrelevant antigen, the 2^nd^ row % Arrested on their cognate antigen and the last row shows the ratio between the positive to negative (1^st^ row to 2^nd^ row). (E) Data showing monoclonal yeast at varying shear rates in a similar format as D.

We tested the Nef19^+^ and CD16.21^+^ yeasts on 6 different concentrations (0.56-135 nM) of their cognate or irrelevant antigen. Conversion of antigen incubation concentration to surface density of antigen on the chamber floor is further discussed in the Suppl. Mat. The general observed tendencies were a higher specific capture fraction at higher antigen concentrations for both antigens (Fig. 2D). However, the nonspecific capture also increased with increasing antigen concentration. The shear rates were varied between 168 and 926 1/s. The lowest tested shear rate of 168 1/s generally led to a high nonspecific capture that dropped to values around 1 % as soon as a shear rate of 337 1/s was used (Fig. 2E). The non-specific capture was generally independent of shear rate, whereas the specific capture decreased markedly with increasing shear rate.

To evaluate the optimal conditions leading the highest specific capture, we calculated the ratios of the capture fraction on the cognate antigen to the capture fraction on the irrelevant antigen. For CD16.21^+^ yeast, the condition with the highest ratio of positive to negative capture fractions was at 45 nM concentrations at a shear rate of 926 1/s, yielding a ratio of 124. For the Nef19^+^ yeast, the condition with the highest ratio was also at 45 nM but at a shear rate of 337 1/s with a ratio of 86. The antigen concentration of 45 nM, equivalent to 180 molecules/µm^2^, thus consistently provided the highest ratio. Overall, the ratios were roughly independent of the shear rate above 168 1/s, being mostly set by the non-specific capture. We decided to move forward with the lower shear rate of 337 1/s, minimizing the shear rate applied while leading to less than 1 % nonspecific capture for both antigens.

We can estimate the number of bonds formed between the Nbs on the yeast and the antigen on the chamber surface. First, considering the length of the stalk between the Nb and the yeast surface (*L*=100 nm) and the typical yeast radius *(a* = 5 µm), the surface of contact between the cell and surface where the ligand and receptor can form a bond is *s* = *π.L.(2a-L)* = 3 µm². The antigen density on the channel surface based on our previous work (17) showed an incubation concentration of 7 nM corresponded to a density 30 molecules/µm². Here, an incubation concentration of 45 nM leads to an estimated antigen density *d_Ag_* = 180 molecules/µm². The maximal number of antigens on the substrate which can form a bond with a Nb on the surface of the yeast in contact with the substrate can then be estimated as: *N_AgContact_* = *d_Ag_. s,* ∼ 500 molecules. A specific yeast display expression vector using Aga2p is known to express around *N_NbTotal_* = 10^4^ – 10^5^ molecules per yeast but can vary between individual cells (45). If we assume similar expression levels, we could therefore estimate the number of Nbs in the contact surface *N_NbContact =_ s.N_NbTotal_ / 4π.a²*, yielding a maximum of 100 Nbs for *N_Nbtotal_* = 10^4^ and a maximum of 1000 Nbs for *N_Nbtotal_* = 10^5^. Thus, the maximal number of bonds *N_max_*, being the minimum of *N_AgContact_* and *N_NbContact_*, should be limited by the number of antigens (500) for yeast with high nanobody display, or by the number of displayed nanobodies (100) for yeast with low nanobody displayWe note that this high number of bonds will favour avidity as the control parameter of selection. To prevent an avidity effect during yeast arrest, the density of antigens on the surface can be highly diluted, so that statistically only one antigen molecule is available at a time for each binding yeast. This is the limit commonly achieved for single bond measurements with the laminar flow chamber (10,17,46). In this limit, the probability of capture is only related to the apparent affinity Kd under flow which can be tuned by the velocity. The total force applied to a yeast can be estimated at *F* = 2 nN using the following formula (22):

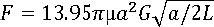

where *µ* = 0.001 Pa.s is the medium viscosity. 1/sFor G = 337 1/s and if the force is equally shared between all the *N_max_* bonds, the force per bond should vary between 4 pN (for highly covered yeast) and 20 pN (for sparsely covered yeast).

### Enrichment using model mixtures of Nb-expressing yeast

Model selections were next performed using the optimized conditions in a new set up equipped with 2 entries -2 exits for better control of the elution step as shown in Fig 3A & 3B, respectively.

**Figure 3.**
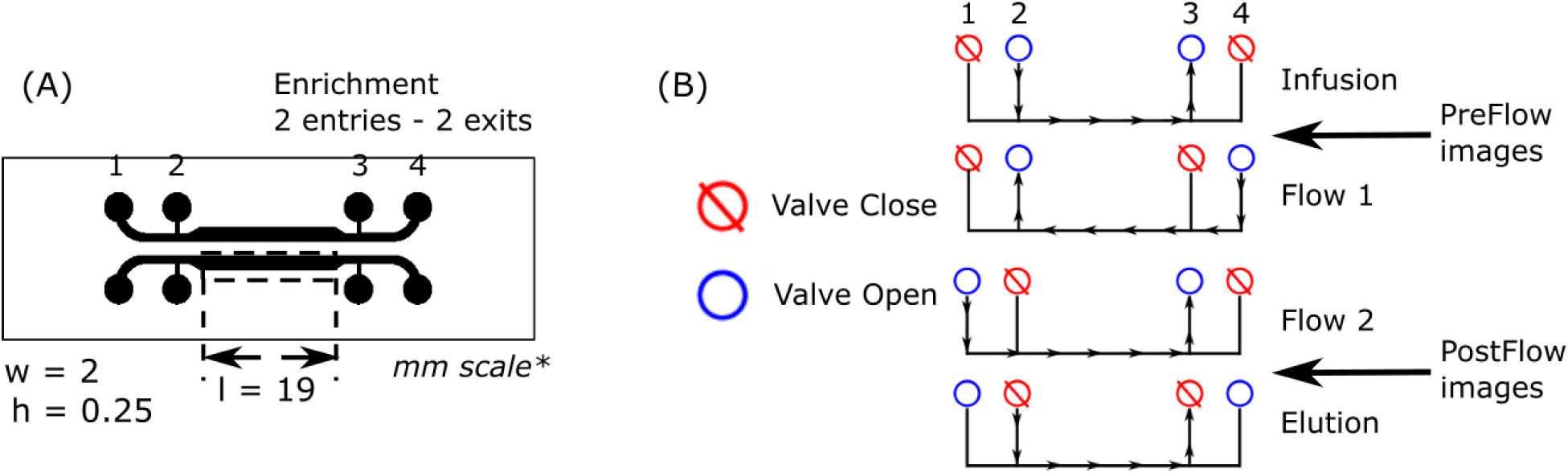
Enrichment Assay microfluidic chamber dimensions & sequence of steps. (A) 2 entries – 2 exits channel used for enrichment based from the design from Rienmets et al. 2019 (37). (B) The 4-step wash sequence used with corresponding closed and opened valves and the direction of flow applied.

To maintain the same shear rate *G* as the previous device, we applied a shear rate of 337 1/s. We used 3 different model mix with ratio of binders to non-binders of 1:1, 1:10 and 1:100. The antigen concentration on the channel surface was maintained at 45 nM for both CD16a-biotin and Nef-biotin. These mixes were subjected to one round of LFC-based enrichment performed at a shear rate of 337 1/s and elution was performed using a high shear rate of 4800 1/s . PreFlow and PostFlow cell counts were recorded and pure population samples were used as reference. As assessed by microscopy (Fig 4A), the percentages of arrested cells for the 1:1 mix (Fig 4B) driven along the Nef-biotin and CD16a-biotin derivatized channel surfaces were at 5 % and 4 % respectively, compared with 2 % and 2.4 % for the 1:10 mixes (Fig 4C), and 1 % for the 1:100 CD16.21^+^: Nef19^+^ mix on the CD16a-biotin channel (Fig 4D). To monitor the enrichment in real time and *in situ*, we stained the non-binding yeast with CFSE to visualize their presence before and after flow for ratios 1:10 and 1:100. A representative image in bright field and corresponding fluorescence is shown in Fig 4A, showing almost no fluorescent cells after flow. Specific and non-specific adhesion values were measured by calculating the ratios of fluorescent cells to non-fluorescent ones before and after flow to obtain the fraction of fluorescent non-binders (% Fluorescent cells). By subtracting this fraction from 1, we get the estimated fraction of non-fluorescent binders (Fig 4E-G).

**Figure 4.**
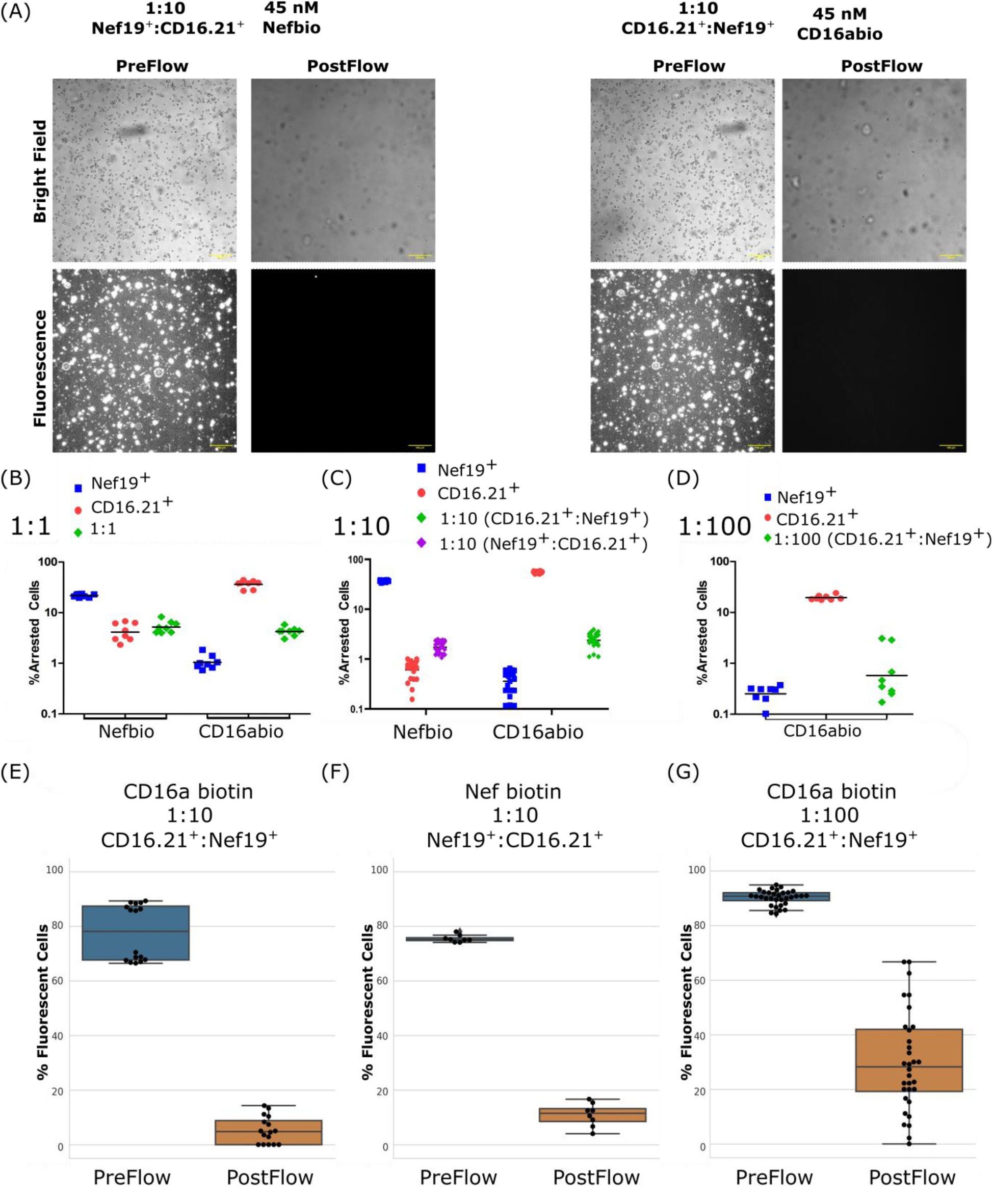
Monitoring of enrichment by microscopy. (A) An example image of the 1:10 mixes used for enrichment. The top row corresponds to the bright field PreFlow & PostFlow images and the lower images are the corresponding fluorescence images. The fluorescent cells are the non-binder cells. The scale bars correspond to 100 µm. (B-D) % Arrested cells of the different ratios (1:1, 1:10 & 1:100) and pure populations (blue and red) measured using bright field, PreFlow and PostFlow. (E-G) % Fluorescent cells of different model mixes before and after flow.

To confirm that enrichment did occur using a different approach, we used again flow cytometry. After the procedure, the eluted cells were amplified and nanobody expression was restored using a 3 days culture, before flow cytometry analysis. Using the quadrant definition shown in Fig1D, we estimated the fraction of positive cells before and after flow using the formula Q2/(Q2+Q3) (see the Suppl. Mat. for the discussion of the limits of this choice) (Fig 5). For the 1:1 ratio, the ratio of binders to expressors increased from 0.49 to 0.66 and 0.86 when driven along the channel surface derivatized with CD16a-biotin and Nef-biotin respectively, i.e., yielding enrichment of 1.3 and 1.8.

**Figure 5.**
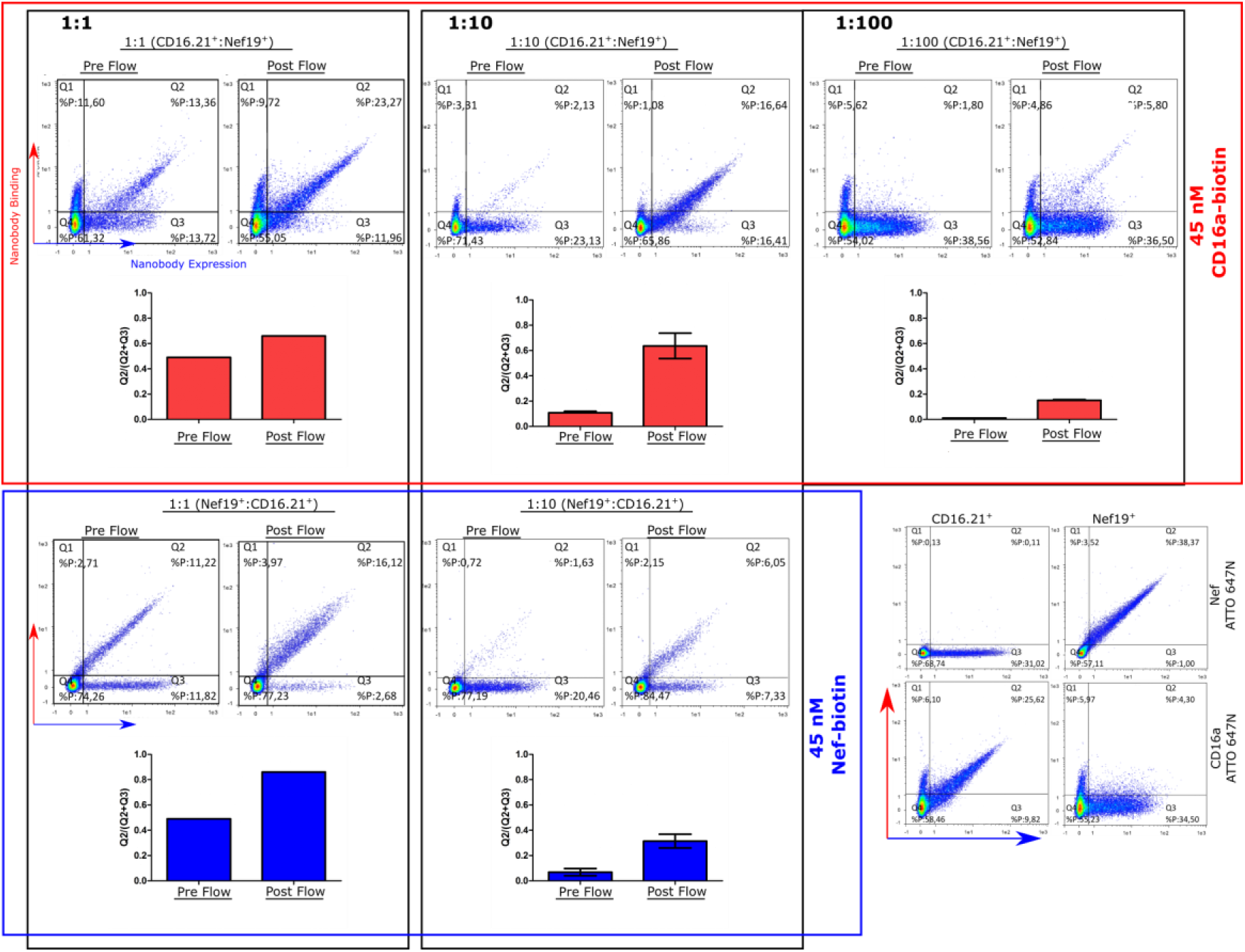
Monitor of Enrichment by Cytometry. The eluted cells were grown and characterized by flow cytometry. The top row shows the cytometry results and the estimated enrichment factor of all CD16.21^+^: Nef19^+^ mixes driven along a CD16a-biotin surface. The bottom row corresponds to the results of all different mixtures of Nef19^+^:CD16.21^+^ driven over a Nef-biotin surface. The right bottom corner displays results obtained with pure populations of CD16.21^+^ or Nef19^+^ yeast with their corresponding cognate or irrelevant antigen as reference.

For the 1:10 CD16.21: Nef19^+^ mix over CD16a-biotin, the ratio of binders to expressors increased from 0.11 to 0.64, an enrichment factor of 5.8. For the 1:10 mix of Nef19^+^:CD16.21^+^ driven along a Nef-biotin functionalized channel surface, this ratio increased from 0.07 to 0.31, i.e., an enrichment factor of 4.4. The 1:100 ratio was only tested on CD16.21^+^: Nef19^+^ mix. Using this ratio, below 1 % of positive cells expected before enrichment falls below the background signal by flow cytometry and thus cannot be measured efficiently. Hence, instead of relying on the direct cytometry data, we used a theoretical value of 0.01 corresponding to the expected 1:100 mixture. The ratio of binders to expressors increased to 0.15, yielding an enrichment factor of 15.

Thus, using this flow-based assay, we were able to reach up to a 15-fold enrichment of positive cells after a single round.

### Predicting the Enrichment from adhesion

To further evaluate these results, we sought to predict the enrichments, as measured using cytometry and based on the adhesion measured in flow chamber by microscopy. We used the following nomenclatures: *f^+^*as the fraction of expressing positive cells, *f^-^* as the fraction of expressing negative cells, *a^+^* as the captured fraction of positive cells after flow, *a^-^* as the captured fraction of negative cells after flow. Assuming that non-expressing positive cells adhere similarly to negative ones (*a^-^*), a theoretical enrichment (*ε*) may be calculated as follows (see the derivation in Suppl. Mat. Eq. (S1)):

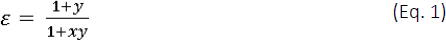

We can define a capture efficiency factor *y* as the product of f^+^ and a term characterizing the selectivity of the channel dictated by the functionalized antigen on its surface

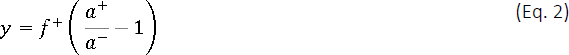

Figure 6A shows the theoretical enrichment *ε* of a model mixture as a function of the initial mix ratio *x* and calculated for various values of capture efficiency *y*. This shows how the enrichment is maximal for low mix ratio *x* but limited by the capture efficiency *y*+1. *y* can be measured using negative and positive adhesion tests or using the fluorescently labelled negative cells (see Suppl. Mat). Using Eq. 1 on our assays lead to a predicted enrichment ε similar for the 2 methods (See Suppl. Mat. Fig 4A, 4B, 4D & 4E). The capture efficiency *y* shows that the order of calculated *y* for the 1:10 ratios using Bright Field and Fluorescence were reversed between the 2 antigens but are both still within the same scale (Suppl. Mat Fig 4C & 4F). Thus, both *in situ* monitoring methods (Bright Field and Fluorescence) could be used to predict enrichment factors (Fig 6B & 6C) that are in good agreement with the enrichments experimentally measured using flow cytometry (Fig 6D) after expansion of the enriched cell populations.

**Figure 6.**
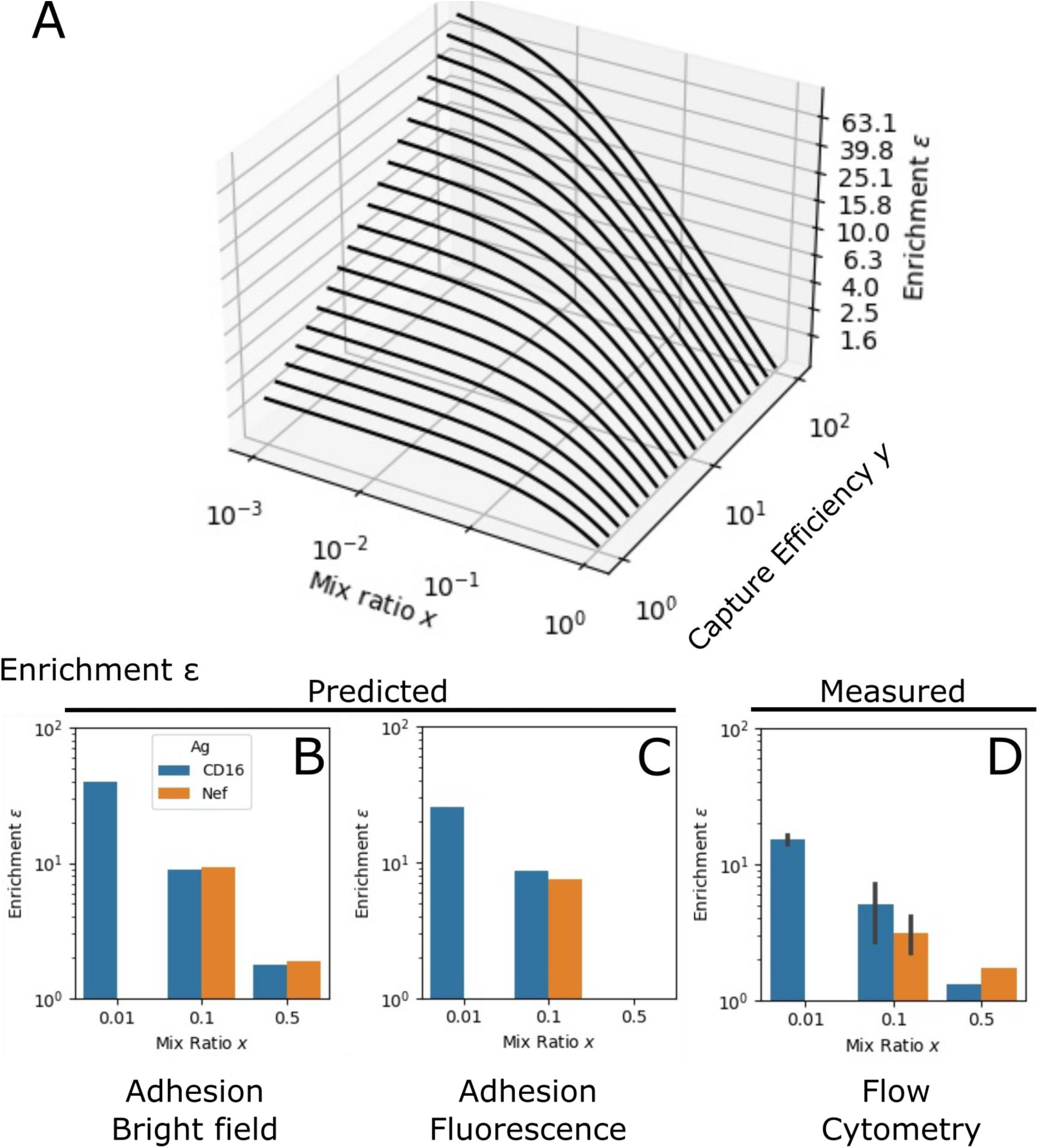
Measured and predicted enrichment of model mixes. (A) Theoretical enrichment *ε* as a function of the mix ratio *x* and Capture Efficiency *y*, using Eq. 1 & Eq 2. (B) The predicted enrichment *ε* based on the % of Arrested cells in bright field microscopy. (C) The predicted enrichment *ε* based on the % Fluorescent Cells in Fluorescence microscopy. (D) Enrichment measured by cytometry PreFlow and PostFlow. The bar graph colors correspond to the immobilized antigen (blue for CD16, orange for Nef).

Applying Eq. 1, one gets a theoretical enrichment *ε* measured through two different microscopy modalities that is consistent with the measured enrichment via cytometry. As a theoretical exercise mimicking a more realistic situation with rare binders within a library, such as a mix ratio *x* of 1:10^6^ and assuming capture efficiency *y* of 50, the fraction of positive yeasts will be 5.0x10^-5^ after round 1, 2.6x10^-3^ after round 2 and 1.3x10^-1^ after round 3, reaching a proportion of binders compatible with random picking of clones for deeper characterization, i.e. a frequency above 10% of positive clones. Of note, reducing the antigen density to avoid avidity effect is expected to yield a lower adhesive ratio. It may thus be applied after a first round of selection at high density, allowing a reduction of diversity and an increase in the copy number of each clone. Alternatively, a preliminary round using magnetic enrichment can also be used to reduce diversity prior to the use of the flow chamber.

## Conclusion

This work validated the efficacy of a microfluidic assay to quantify both the antigen-specific and non-specific capture fraction of yeasts expressing a specific Nb on their surface and translating it into a method for enrichment of a model mix. This result was achieved through the precise control of shear rates and antigen surface densities, and the use of precise washing and elution steps. Our assay enabled the detection of antigen-specific capture and enrichment for yeasts that express antigen-specific Nbs.

Importantly, our method provides a way to control the force applied to the interaction by modulating the shear rate, as well as the valency of the interaction, by modulating the antigen surface density. These are major parameters to ultimately tune the selection of binders with pre-determined force response and single/multivalent bond behaviour. Current yeast display strategies mainly use magnetic activated cell sorting or fluorescence activated cell sorting, two methods relying on affinity that do not take into consideration the forces surrounding the receptor-ligand interaction. Conversely, our force-based selection strategy has the potential to favor the selection of binders able to withstand a certain amount of external force. By allowing the force-based enrichment of yeasts displaying libraries of antibody fragments such as nanobodies, scFv, Fab, but also full-length antibodies, this strategy may represent an important step toward the engineering of more efficient immunotherapeutics.

## Author Contributions

M. S. performed the experiments and analysed the data. M.S. was responsible for all cell manipulations, microfluidic channel fabrications and imaging. R.T. developed the image analysis software. L.L. derived the theoretical prediction model for enrichment. All authors conceived and designed the experiments and wrote and edited the manuscript.

## Acknowledgements

We thank Yong Jian Wang, Dalia El Arawi, Luc David-Bruglio and Pierre Bohec for technical help with the microfluidics as well as Philippe Robert and Olivier Theodoly for fruitful discussions. We also thank Elise Termine, Remi Bonjean and Adrian Aimard for technical help with protein production and labelling and Timothee Chanier for technical help with flow cytometry. This work has been carried out thanks to the support of the A*MIDEX project (ANR-11-IDEX-0001-02) funded by the Investissements d’Avenir program from the French government, managed by the French National Research Agency (ANR); and of the Physcancer program from Institut National du Cancer-Plan Cancer.

## Conflict of Interest

There are no conflicts to declare.

## Supplementary Materials

### Antigen Production and Conjugation

Nef-biotin was produced using BL21DE3 *E. coli* co-transformed with Nef-AviTag-6His and BirA-cm via heat shock. Transformed cells were grown in a 5 mL 2YT medium (16 g Tryptone, 10 g Yeast Extract, 5 g NaCl with 1 L MilliQ water) supplemented with 2 % v/v glucose, 100 µg/mL ampicillin and 50 µg/mL chloramphenicol and incubated at 37 °C shaking at 220 rpm for 5 hours. An appropriate volume from this starter was transferred into 100 mL of a similar media but without the 2 % v/v glucose to have a starting OD_600nm_ = 0.1. The culture was grown until an OD_600nm_ = 0.5-0.8 was reached and induced by adding 100 µM Isopropyl β-D-1-thiogalactopyranoside (IPTG) and 10 µM biotin and incubated overnight at 30 °C. Afterwards the cells were lysed using a mixture of BugBuster with Benzonase and Lysozine purified using cobalt resin (TALON superflow, GE Healthcare).

Nef-cmyc was produced using BL21DE3 *E. coli* transformed with Nef-cmyc-6His via heat shock. Transformed cells were grown in a 5 mL 2YT medium supplemented with 2 % v/v glucose and 100 µg/mL ampicillin and incubated at 37 °C shaking at 220 rpm for 5 hours. An appropriate volume from this starter was transferred into 100 mL of a similar media but without the 2 % v/v glucose to have a starting OD_600nm_ = 0.1. The culture was grown until an OD_600nm_ = 0.5-0.8 was reached and induced by adding 100 µM Isopropyl β-D-1-thiogalactopyranoside (IPTG) and incubated overnight at 30 °C. Afterwards the cells were lysed and purified as previously described.

CD16a-cmyc was produced using Expi293F cells (A14635, Gibco^TM^) transformed with CD16a-cmyc-6His using Expifectamine DNA lipid complex as described in the product notes. After 1 week of growth, the supernatant was recovered and underwent overnight dialysis using Spectra/Por® 4 RC Dialysis Membrane Tubing at 2 mL/cm with a MWCO of 12,000-14,000 kD in PBS 1x. Purification was also done using cobalt resin.

Nef-cmyc and a portion of CD16a-cmyc underwent labelling with ATTO 647N using the bacterial transglutaminase (L107, TGase Q Protein Labeling Kit, Zedira) as described in the product notes. A portion of the CD16a-cmyc underwent conjugation with biotin using bacterial transglutaminase (L101) as described in the product notes.

### CFSE Staining of Negative Yeast

CFSE (CellTrace, C34554A) staining of yeast was adapted from the supplier provided notes and from this staining protocol (41). Stock CFSE was reconstituted with 18 µL DMSO to create a starting concentration of 5 mM. The staining was done on induced yeasts washed and prepared to have an OD_600nm_ = 1 in 1x PBS by adding CFSE stock at a ratio of 1:1000 to reach a working concentration of 5 µM. This was incubated at room temperature on a Stuart tube rotator at 40 rpm for 30 min. Afterwards the yeast was washed twice; this was done by centrifuging the sample at 3500 x g for 1 min at room temperature and resuspending in PBS 1x with 2 % BSA. Staining was checked using cytometry and microscopy. The staining was generally performed on model enrichment trials with non-binding yeasts stained with CFSE, excluding the 1:1 model mixture.

### Estimation of Antigen Concentration on the channel surface

In a previous publication (17), it was shown in a microfluidic chamber with glass bottom and PDMS channel, a 7 nM incubation concentration yielded a surface concentration of 30 molecules/µm^2^, a conversion factor of 4. Applying this to the concentration used for enrichment, the 45 nM incubation concentration will have 180 molecules/µm^2^. For the serial dilution in the optimization of the incubation concentration to be used in the microfluidic channel, the 135, 45, 15, 5, 1.67 and 0.56 nM converts to 540, 180, 60, 20, 7 and 2 molecules/µm^2^, respectively.

### Derivation of Eq. (1) and (2) from the main text

*E^+^* = fraction of Expressor Positive (Binding) Cells

*E^-^* = fraction of Expressor Negative (Non-binding) Cells

*NE^+^* = fraction of Non-Expressor Positive Cells (contains the plasmid but not expressing the Nb)

*NE^-^* = fraction of Non-Expressor Negative Cells

*f^+^* = *E^+^/ (E^+^ + NE^+^)* = fraction of Expressing Positive Cells

*f^-^* = *E^-^/ (E^-^ + NE^-^)* = fraction of Expressing Negative Cells

*a^+^*= captured fraction of Positive Cells after flow

*a^-^*= captured fraction of Negative Cells after flow

The sum of *E^+^, NE^+^, E^-^* & *NE^-^* equals 1. We assume x as the ratio of positive cells (bearing the positive plasmid) before flow where *x = E^+^_pre_ + NE^+^_pre_*. Therefore, the fractions can be expressed as in Table 1.

**Table 1.**
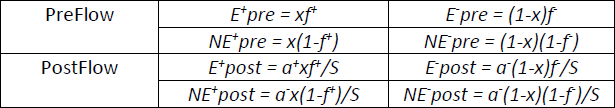
Fractions PreFlow and PostFlow.

A sum *S* was applied to normalize the data and ensure a 1 for the sum of all fractions PostFlow. Additionally, we assume that non-expressing positive cells (*NE^+^*) adhere like negative ones (*a^-^*).

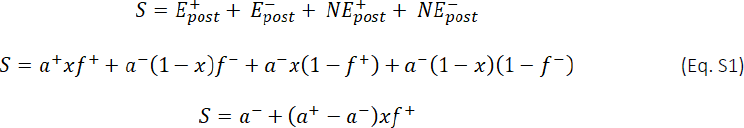

A theoretical enrichment (*ε*) in positive cells is defined as:

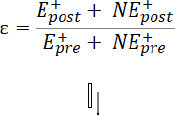

Adding the mix ratio x and the equations from Table 1, we get the following:

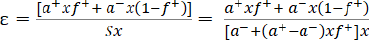

leading to Eq. 1 and 2 in the main text.

### Measurement of *y* using pure yeast population

The fraction of cells adhered after flow for a pure yeast population with the irrelevant Nb is: (*N_post_*/*N_pre_*)^-^= *S*(*x*=0) =*a^-^*.

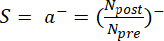

On the other hand, the fraction of cells adhered for a pure yeast population with the cognate Nb is:

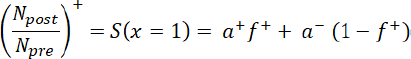

When we replace these quantities from the previous equation of *y* we get:

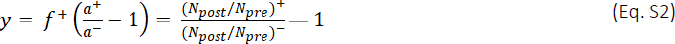

Interestingly, this equation shows that the measured capture efficiencies using the monoclonal control populations driven along a surface with an antigen of interest may be used to predict the theoretical enrichment *ε*.

### Measurement of *y* using fluorescent negative yeasts in the mixture

Alternatively, the data from the *in-situ* fluorescence microscopy can be used to measure the adhesion. The number of positive cells corresponds to the total number of cells observed in bright field (BF) minus the number of fluorescent ones (Fluo). One defines therefore the negative and positive capture efficiencies respectively as:

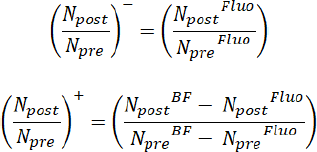

which are used in Eq. S2 to determine *y*.

### Cytometry Correction for Enrichment Prediction

In an ideal situation, Q1 would not contain any signal. So, if we assume *f^+^* = *f^-^*, and Q1 = 0, we can calculate *E^+^* = Q2, *E^-^* = Q3, *NE^+^*= Q2Q4/(Q2+Q3) and *NE^-^* = Q3Q4/(Q2+Q3). In this case, we can calculate the mix ratio *x* as well as the enrichment ε as a function of Q2, Q3 and Q4. However, cytometry data show signals for both Q1 and Q2 even on a negative control. Using control yeasts in cytometry, we computed for autofluorescence (*f_AF_*) and obtained a corrected *f^+^*as shown in the table below.

**Table 2.**
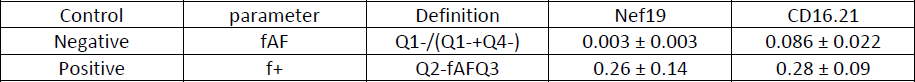
Autofluoresence f_AF_ and expression fraction f^+^ established using Controls.

The estimate for enrichment in the in-text Fig 5 was through Q2/ (Q2+Q3) for PreFlow and PostFlow which represents the fraction of binders over expressors and considering the ratio PostFlow/PreFlow. We can calculate those quantities using fractions, Q2 = *E^+^* and Q3 = *E^-^* and assume that *f^+^* = *f^-^* = *f*.

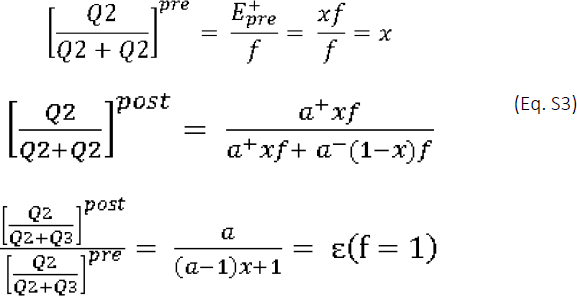

This is only applicable in a condition where *f* = 1, a condition that is not fulfilled as seen in Table 2. Thus, the ratio Q2/ (Q2+Q3) roughly underestimated the real enrichment factor if cytometry would be performed immediately after the selection. In practice, the cytometry measurements PostFlow is performed after 48-72h of cell culture to allow for cell expansion and induction of expression, a time sufficient to recover the initial fraction of expressors *f^+^*. We thus considered the enrichment estimated from Q2/ (Q2+Q3) to be valid. The predicted enrichment *ε*(*f*=1) can also be calculated as the following: The factor *y*(*f^+^*) which corresponds to the real expression fraction *f^+^* is obtained using the adhesion measurements. We deduced the value of y corresponding to *f* = 1: *y*(*f*=1) = *y(f^+^)/f^+^* therefore:

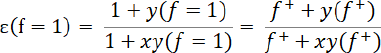

**Supplementary Material Figure 1.**
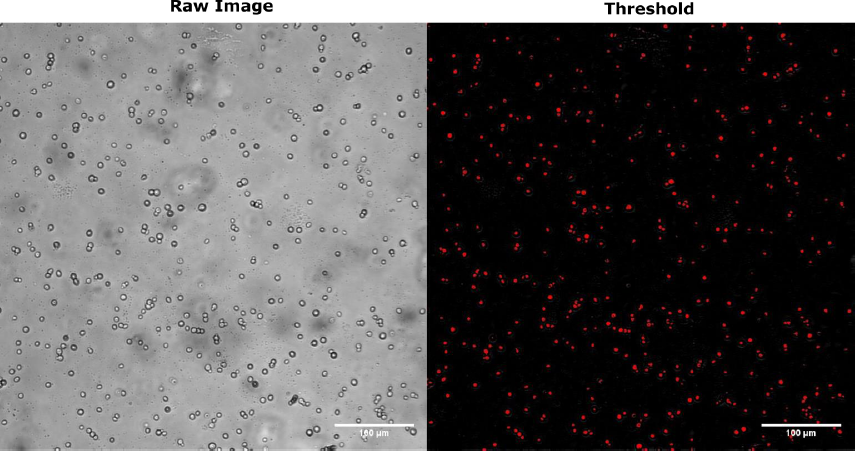
Detection of Yeast Cells. Detection and counting of yeast cells was done using the MorphoLibJ plugin function Gray Scale Attribute Filtering with Top Hat in FIJI v1.53t. Here we show an example of an image in BF microscopy and the corresponding image after thresholding. The red dots on black background show the yeast cells detected and counted. White Scale bar is at 100 µm.

**Supplementary Material Figure 2.**
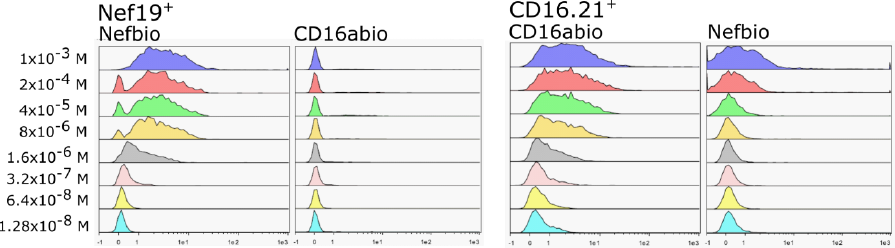
Cytometry Histograms to estimate apparent affinity. The apparent affinity of the Nb on the yeast surface was estimated. Nef19^+^ & CD16.21^+^ were incubated with either their cognate or irrelevant antigen and differing concentrations. Each concentrations used are indicated on the corresponding row. Values used in main text Fig 1C were taken here gated using the lowest concentration on the irrelevant antigen.

**Supplementary Material Figure 3.**
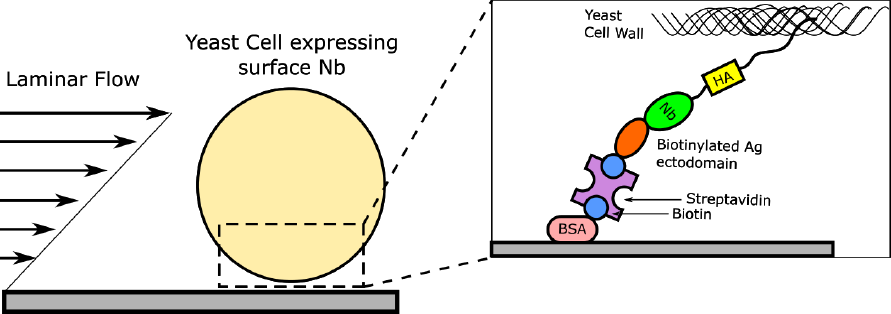
Yeast driven along the channel surface. A schematic showing the expected interaction between the flowing yeast cell on the channel surface and the antigen functionalized on the channel surface (image not to scale).

**Supplementary Material Figure 4.**
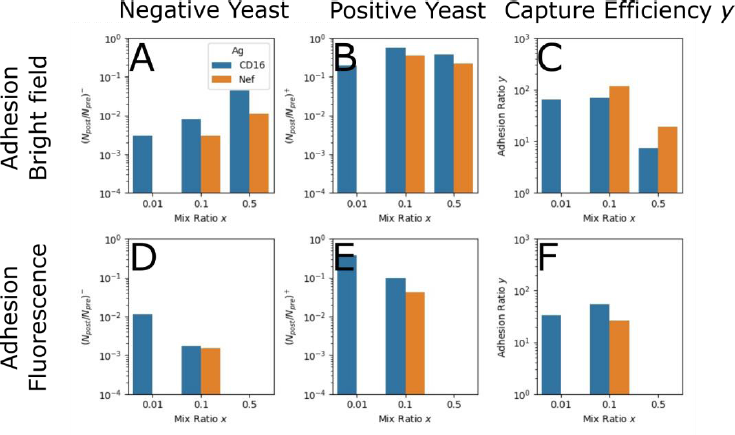
Bright Field (BF), Fluorescence (Fluo) and Capture Efficiency *y*. (A) The fraction of cell count PostFlow to PreFlow of the pure non-binding yeast (negative) used as control during enrichment experiments imaged through BF Microscopy. (B) The fraction of cell count PostFlow to PreFlow of the pure binding yeast (positive) used as control during enrichment experiments imaged through BF Microscopy. (C) The calculated capture efficiency *y* using the BF data using Eq 2 of the main text. (D) The fraction of cell count PostFlow to PreFlow of the pure non-binding & fluorescent yeasts (negative) in the mixture during enrichment experiments imaged through Fluo Microscopy. (E) The fraction of cell count PostFlow to PreFlow of the pure non-binding & non-fluorescent yeast (positive) in the mixture during enrichment experiments imaged through Fluo Microscopy. (F) The calculated capture efficiency *y* using the Fluo data using Eq 2 of the main text.

## Notes

### Competing Interest Statement

The authors have declared no competing interest.

## References

1. Huse M. Mechanical Forces in the Immune System. Nat Rev Immunol. 2017 Nov;17(11):679–90.

2. 2. Kopf A, Kiermaier E. Dynamic Microtubule Arrays in Leukocytes and Their Role in Cell Migration and Immune Synapse Formation. Front Cell Dev Biol [Internet]. 2021 [cited 2023 Feb 2];9. Available from: https://www.frontiersin.org/articles/10.3389/fcell.2021.635511

3. Dustin ML, Chakraborty AK, Shaw AS. Understanding the Structure and Function of the Immunological Synapse. Cold Spring Harb Perspect Biol. 2010 Oct 1;2(10):a002311.

4. Bashour KT, Gondarenko A, Chen H, Shen K, Liu X, Huse M, et al. CD28 and CD3 have complementary roles in T-cell traction forces. Proc Natl Acad Sci. 2014 Feb 11;111(6):2241–6.

5. Göhring J, Kellner F, Schrangl L, Platzer R, Klotzsch E, Stockinger H, et al. Temporal analysis of T-cell receptor-imposed forces via quantitative single molecule FRET measurements. Nat Commun. 2021 May 4;12(1):2502.

6. Ganti RS, Lo WL, McAffee DB, Groves JT, Weiss A, Chakraborty AK. How the T cell signaling network processes information to discriminate between self and agonist ligands. Proc Natl Acad Sci. 2020 Oct 20;117(42):26020–30.

7. Hong J, Persaud SP, Horvath S, Allen PM, Evavold BD, Zhu C. Force-Regulated In Situ TCR– Peptide-Bound MHC Class II Kinetics Determine Functions of CD4+ T Cells. J Immunol. 2015 Oct 15;195(8):3557–64.

8. Wang J huai. T cell receptors, mechanosensors, catch bonds and immunotherapy. Prog Biophys Mol Biol. 2020 Jul 1;153:23–7.

9. Sibener LV, Fernandes RA, Kolawole EM, Carbone CB, Liu F, McAffee D, et al. Isolation of a Structural Mechanism for Uncoupling T Cell Receptor Signaling from Peptide-MHC Binding. Cell. 2018 Jul;174(3):672–687.e27.

10. Limozin L, Bridge M, Bongrand P, Dushek O, van der Merwe PA, Robert P. TCR–pMHC kinetics under force in a cell-free system show no intrinsic catch bond, but a minimal encounter duration before binding. Proc Natl Acad Sci. 2019 Aug 20;116(34):16943–8.

11. Le Saux G, Bar-Hanin N, Edri A, Hadad U, Porgador A, Schvartzman M. Nanoscale Mechanosensing of Natural Killer Cells is Revealed by Antigen-Functionalized Nanowires. Adv Mater. 2019;31(4):1805954.

12. Fan J, Shi J, Zhang Y, Liu J, An C, Zhu H, et al. NKG2D discriminates diverse ligands through selectively mechano-regulated ligand conformational changes. EMBO J. 2022 Jan 17;41(2):e107739.

13. Natkanski E, Lee WY, Mistry B, Casal A, Molloy JE, Tolar P. B Cells Use Mechanical Energy to Discriminate Antigen Affinities. Science. 2013 Jun 28;340(6140):1587–90.

14. Jiang H, Wang S. Immune cells use active tugging forces to distinguish affinity and accelerate evolution. Proc Natl Acad Sci. 2023 Mar 14;120(11):e2213067120.

15. Faro J, Castro M. Affinity Selection in Germinal Centers: Cautionary Tales and New Opportunities. Cells. 2021 May;10(5):1040.

16. Hamers-Casterman C, Atarhouch T, Muyldermans S, Robinson G, Hammers C, Songa EB, et al. Naturally occurring antibodies devoid of light chains. Nature. 1993 Jun;363(6428):446–8.

17. González C, Chames P, Kerfelec B, Baty D, Robert P, Limozin L. Nanobody-CD16 Catch Bond Reveals NK Cell Mechanosensitivity. Biophys J. 2019 Apr 23;116(8):1516–26.

18. Heath GR, Kots E, Robertson JL, Lansky S, Khelashvili G, Weinstein H, et al. Localization atomic force microscopy. Nature. 2021 Jun;594(7863):385–90.

19. Kostrz D, Wayment-Steele HK, Wang J, Follenfant M, Pande VS, Strick TR, et al. A modular DNA scaffold to study protein protein interactions at single-molecule resolution. Nat Nanotechnol. 2019;14(10):988.

20. Sitters G, Kamsma D, Thalhammer G, Ritsch-Marte M, Peterman EJG, Wuite GJL. Acoustic force spectroscopy. Nat Methods. 2015 Jan;12(1):47–50.

21. Wang YJ, Valotteau C, Aimard A, Villanueva L, Kostrz D, Follenfant M, et al. Combining DNA scaffolds and acoustic force spectroscopy to characterize individual protein bonds. bioRxiv. Accepted to Biophysical Journal.; 2022. p. 2022.08.14.503897.

22. Pierres A, Benoliel AM, Bongrand P. Measuring the Lifetime of Bonds Made between Surface-linked Molecules (∗). J Biol Chem. 1995 Nov 3;270(44):26586–92.

23. Pierres A, Feracci H, Delmas V, Benoliel AM, Thiery JP, Bongrand P. Experimental study of the interaction range and association rate of surface-attached cadherin 11. Proc Natl Acad Sci. 1998 Aug 4;95(16):9256–61.

24. Limozin L, Bongrand P, Robert P. A Rough Energy Landscape to Describe Surface-Linked Antibody and Antigen Bond Formation. Sci Rep. 2016 Oct 12;6(1):35193.

25. Li P, Gao Y, Pappas D. Multiparameter Cell Affinity Chromatography: Separation and Analysis in a Single Microfluidic Channel. Anal Chem. 2012 Oct 2;84(19):8140–8.

26. Cheng X, Irimia D, Dixon M, Sekine K, Demirci U, Zamir L, et al. A microfluidic device for practical label-free CD4+ T cell counting of HIV-infected subjects. Lab Chip. 2007;7(2):170–8.

27. Pullagurla SR, Witek MA, Jackson JM, Lindell MAM, Hupert ML, Nesterova IV, et al. Parallel Affinity-Based Isolation of Leukocyte Subsets Using Microfluidics: Application for Stroke Diagnosis. Anal Chem. 2014 Apr 15;86(8):4058–65.

28. Smith GP. Filamentous Fusion Phage: Novel Expression Vectors That Display Cloned Antigens on the Virion Surface. Science. 1985 Jun 14;228(4705):1315–7.

29. Boder ET, Wittrup KD. Yeast surface display for screening combinatorial polypeptide libraries. Nat Biotechnol. 1997 Jun;15(6):553–7.

30. Oliphant T, Engle M, Nybakken GE, Doane C, Johnson S, Huang L, et al. Development of a humanized monoclonal antibody with therapeutic potential against West Nile virus. Nat Med. 2005 May;11(5):522–30.

31. Kalb SR, Lou J, Garcia-Rodriguez C, Geren IN, Smith TJ, Moura H, et al. Extraction and Inhibition of Enzymatic Activity of Botulinum Neurotoxins/A1, /A2, and /A3 by a Panel of Monoclonal Anti-BoNT/A Antibodies. PLoS ONE. 2009 Apr 28;4(4):e5355.

32. McMahon C, Baier AS, Pascolutti R, Wegrecki M, Zheng S, Ong JX, et al. Yeast surface display platform for rapid discovery of conformationally selective nanobodies. Nat Struct Mol Biol. 2018 Mar;25(3):289–96.

33. Pymm P, Redmond SJ, Dolezal O, Mordant F, Lopez E, Cooney JP, et al. Biparatopic nanobodies targeting the receptor binding domain efficiently neutralize SARS-CoV-2. iScience. 2022 Nov 18;25(11):105259.

34. Teymennet-Ramírez KV, Martínez-Morales F, Trejo-Hernández MR. Yeast Surface Display System: Strategies for Improvement and Biotechnological Applications. Front Bioeng Biotechnol. 2022 Jan 10;9:794742.

35. Bowley DR, Labrijn AF, Zwick MB, Burton DR. Antigen selection from an HIV-1 immune antibody library displayed on yeast yields many novel antibodies compared to selection from the same library displayed on phage. Protein Eng Des Sel. 2007 Jan 1;20(2):81–90.

36. Richter F, Bindschedler S, Calonne-Salmon M, Declerck S, Junier P, Stanley CE. Fungi-on-a-Chip: microfluidic platforms for single-cell studies on fungi. FEMS Microbiol Rev. 2022 Nov 1;46(6):fuac039.

37. Reinmets K, Dehkharghani A, Guasto JS, Fuchs SM. Microfluidic quantification and separation of yeast based on surface adhesion. Lab Chip. 2019 Oct 9;19(20):3481–9.

38. Bouchet J, Basmaciogullari SE, Chrobak P, Stolp B, Bouchard N, Fackler OT, et al. Inhibition of the Nef regulatory protein of HIV-1 by a single-domain antibody. Blood. 2011 Mar 31;117(13):3559–68.

39. Behar G, Sibéril S, Groulet A, Chames P, Pugnière M, Boix C, et al. Isolation and characterization of anti-FcγRIII (CD16) llama single-domain antibodies that activate natural killer cells. Protein Eng Des Sel. 2008 Jan 1;21(1):1–10.

40. Gietz RD, Woods RA. Transformation of yeast by lithium acetate/single-stranded carrier DNA/polyethylene glycol method - ScienceDirect [Internet]. 2002 [cited 2022 Nov 9]. Available from: https://www-sciencedirect-com.lama.univ-amu.fr/science/article/abs/pii/S0076687902509575?via%3Dihub

41. Dagher Z, Xu S, Negoro PE, Khan NS, Feldman MB, Reedy JL, et al. Fluorescent Tracking of Yeast Division Clarifies the Essential Role of Spleen Tyrosine Kinase in the Intracellular Control of Candida glabrata in Macrophages. Front Immunol. 2018 May 16;9:1058.

42. Batchelor GK. An Introduction to Fluid Dynamics. Cambridge University Press; 2000. 615 p.

43. Crocker JC, Grier DG. Methods of Digital Video Microscopy for Colloidal Studies. J Colloid Interface Sci. 1996 Apr 15;179(1):298–310.

44. Kajiwara K, Aoki W, Ueda M. Evaluation of the yeast surface display system for screening of functional nanobodies. AMB Express. 2020 Mar 16;10(1):51.

45. Linciano S, Pluda S, Bacchin A, Angelini A. Molecular evolution of peptides by yeast surface display technology. MedChemComm. 2019 Sep 18;10(9):1569–80.

46. Robert P, Nicolas A, Aranda-Espinoza S, Bongrand P, Limozin L. Minimal Encounter Time and Separation Determine Ligand-Receptor Binding in Cell Adhesion. Biophys J. 2011 Jun 8;100(11):2642–51.

